# Dynamics and Control of Infections on Social Networks of Population Types

**DOI:** 10.1101/200063

**Authors:** Brian G. Williams, Christopher Dye

## Abstract

Random mixing in host populations has been a convenient simplifying assumption in the study of epidemics, but neglects important differences in contact rates within and between population groups. For HIV/AIDS, the assumption of random mixing is inappropriate for epidemics that are concentrated in groups of people at high risk, including female sex workers (FSW) and their male clients (MCF), injecting drug users (IDU) and men who have sex with men (MSM). To find out who transmits infection to whom and how that affects the spread and containment of infection remains a major empirical challenge in the epidemiology of HIV/AIDS. Here we develop a technique, based on the routine sampling of infection in linked population groups (a social network of population types), which shows how an HIV/AIDS epidemic in Can Tho Province of Vietnam began in FSW, was propagated mainly by IDU, and ultimately generated most cases among the female partners of MCF (FPM). Calculation of the case reproduction numbers within and between groups, and for the whole network, provides insights into control that cannot be deduced simply from observations on the prevalence of infection. Specifically, the *per capita* rate of HIV transmission was highest from FSW to MCF, and most HIV infections occurred in FPM, but the number of infections in the whole network is best reduced by interrupting transmission to and from IDU. This analysis can be used to guide HIV/AIDS interventions using needle and syringe exchange, condom distribution and antiretroviral therapy. The method requires only routine data and could be applied to infections in other populations.

## 1. Introduction

Epidemiological theory assumes that infections are transmitted through random contacts between infected and uninfected people. The reality is usually different, and simple assumptions can give misleading results. One example is the spread of HIV/AIDS in ‘concentrated epidemics’, where populations contain small groups of people at high risk and large groups of people at low risk. Various approaches have been developed to analyse and interpret transmission on social networks of population types in which individuals may belong to several different types. In the case under consideration here these types consist of men who have sex with men, MSM, intravenous drug users, IDUs, female sex workers, FSWs, male-clients of FSW, MCF, female partners of MCF that we shall refer to as low risk women, LRW. If a person has only one risk factor, then that determines their population type. If people have more than one risk factor this defines a separate type and the ones of interest here are MSM who are also IDUs, and FSW who are also IDUs giving a total of seven types. It is clear that the population size of the different types, the transmission rate between people of a given type, and the transmission rates between people of different types will vary greatly. Given the differential equations for a model such as this, the calculation of the overall reproduction number, *R*_0_, is straightforward (Diekmann et al., 1991; Diekmann et al., 1990; Diekmann et al., 2010; Heesterbeek, 2002; Roberts and Heesterbeek, 2007). If overall transmission of a pathogen can be reduced by a factor of 1/*R*_0_ then elimination is guaranteed but when the size of groups, the prevalence of the pathogen within each group, their interactions and their risks of infection vary by orders of magnitude, *R*_0_, averaged over the whole network, may not be the most useful guide to controlling the epidemic. To address this issue Heesterbeek and others (Heesterbeek et al., 2015; Heesterbeek and Roberts, 2007; Roberts and Heesterbeek, 2003, 2007, 2012; Shuai et al., 2013) have introduced the Type Reproduction Number *T*_0_. By analogy with *R*_0_, *T*_0_ is the number of secondary cases that arise when one individual of a given population type is introduced into a fully susceptible population of all types. In the case under consideration here, introducing infected one infected FSW will lead to infections in many clients but introducing one infected LRW will lead to no further infections since we assume that they are an epidemiological dead end (Kato et al., 2013). While one would need to calculate *T*_0_ for each of the seven Population Types in the Can Tho network, this provides more nuanced information concerning the optimal strategy for controlling the epidemic. This concept has been further refined by introducing the Target Reproduction Number in which control is targeted at particular interactions between types (Shuai et al., 2013). In Can Tho, for example, male circumcision would reduce transmission from FSW to MCF but not from MCF to FSW. The analysis presented here is essentially an application of the Type Reproduction Number. We consider a situation in which members of a given Type are rendered uninfectious by providing them with anti-retroviral therapy (ART). Given that ART will be rolled-out over time one wishes to determine which Types should be given priority and in what order so as to have the greatest impact on the overall value of *R*_0_. Each time a person is started on ART the number of people of that Type is reduced by 1.

Here we show that, when investigating the control of such epidemics, routinely collected data are a rich source of information. Using surveillance data to characterize the transmission network for HIV/AIDS in Vietnam we find that the best way to minimize infections in the whole population is first by targeting high-risk injection drug users, then men who have sex with men, and finally female sex workers.

Generalized epidemics of HIV/AIDS, such as those prevailing in Eastern and Southern Africa, are driven mainly by heterosexual transmission in the population at large (Gouws and Cuchi, 2012; Williams *et al.*, 2015). Concentrated epidemics, on the other hand, are focused on networked groups of people who acquire and transmit virus by a mix of sexual transmission (between men and women and among men) and by non-medical needle injection of contaminated blood. Investigations of the structure of these networks have usually been carried out with social surveys (Helleringer *et al.*, 2009; Lurie *et al.*, 2003) or by identifying transmission links with genetic markers (Brenner and Wainberg, 2013; Grabowski *et al.*, 2014; Leventhal *et al.*, 2012; Stadler *et al.*, 2012) in order to track the spread of infection through populations and models have been developed to take various levels of network structure into account (Sattenspiel and Simon, 1988). However, the accurate reconstruction of transmission pathways by these methods is labour intensive both in the field and in the laboratory. In this paper we consider an alternative method of constructing an epidemic network based on the routine sampling, through time, of infection in linked population groups. We have used the method to gain insights into the way an epidemic of HIV/AIDS unfolded in Vietnam, and to investigate how the spread of infection can most effectively be reversed.

The control of HIV in concentrated epidemics demands different interventions for different risk groups. In Thailand, the ‘100% Condom Programme’ for female sex workers, combined with other interventions, significantly reduced HIV transmission (Park *et al.*, 2010). For injecting drug users a meta-analyses suggests that access to clean needles and syringes could reduce HIV transmission by 66% (Aspinall *et al.*, 2014) while another meta-analysis suggests the opiate substitution therapy could reduce transmission by 54% (MacArthur *et al.*, 2012). In generalized epidemic settings early treatment has been found to reduce transmission by 96% (Cohen *et al.*, 2011; Cohen *et al.*, 2012). While both the impact and the cost of different combinations of interventions vary, we are concerned in this paper with the population impact that can be achieved for a given reduction in the individual risk of transmission however it is brought about.

## 2. Methods and data

This analysis focuses on the spread of an HIV/AIDS epidemic in Can Tho province, Vietnam, as described by data collected as part of the annual National Sentinel Surveillance system (1994 to 2010) and from Integrated Biological and Behavioural Surveillance surveys in 2006 and 2009. The data used in this analysis, details of the model, choice of parameters and the fitting process are discussed in detail in a previous study (Kato *et al.*, 2013).

In 2010, the prevalence of HIV was highest among injection drug users (IDU: 48%), then men who have sex with men (MSM: 9.5%), followed by female sex workers (FSW: 5.8%), male clients of FSW (MCF: 1.1%) and finally female partners of men in each group (FPM: 0.5%). While the prevalence of infection is lowest in FPM, this group carries the largest number of infections, making up 49% of all infected people, because they are by far the largest group among those at risk of infection.

We use a previously constructed network including transmission within groups and all probable links between pairs of groups (Figure 1) (Kato *et al.*, 2013). Injecting drug users (IDU), men who have sex with men (MSM), and female sex workers (FSW) and their male clients (MCF), each have potentially self-sustaining epidemics. They are connected through MSM and FSW who are also IDU. The female partners of men who visit sex workers (FPM) and of other men are assumed to be an epidemiological dead end, and do not infect anyone else (Kato et *al.*, 2013). In Figure 1, the weight of the arrows indicates the expected extent of transmission. For example, each FSW may infect many MCFs but each MCF is likely to infect relatively few FSWs.

**Figure 1.**
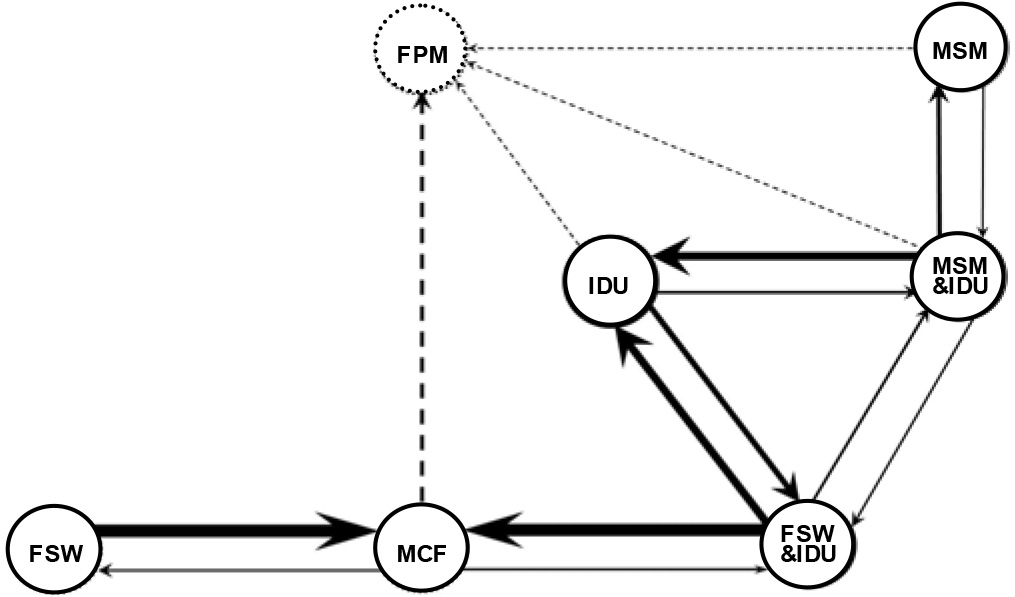
The network model for HIV in Can Tho Province, Viet Nam. IDU: Injection drug users; MSM: Men who have sex with men; FSW: Female sex workers; MCF: Male clients of FSWs; FPM: Female partners of MCF and other women at low risk

The differential equations for the network in Figure 1, are given in the Appendix. The initial prevalence (in 1980) and the transmission parameters were varied to obtain the maximum likelihood fit to the trend data assuming binomial errors (Kato *et al.*, 2013). This gives the estimated size and prevalence in each group and sub-group in 2011 (Table 1) and the fitted trends shown in (Figure 2).

**Table 1.** Risk groups, the estimated number in each group, the prevalence of infection, the number of infected people in each group, and the mean time for which people in high risk groups continue to practice high risk behaviour (Kato et al., 2013). IDU: Injection drug users; MSM: Men who have sex with men; FSW: Female sex workers; MCF: Male clients of FSWs; FPM: Female partners of MCF and other women at low risk.

| Risk group | Number | Prevalence (%) | No. infected | Duration (yrs) |
| --- | --- | --- | --- | --- |
| IDU | 2,716 | 49.50 | 1100 | 12 |
| MSM | 1,176 | 3.62 | 43 | 20 |
| MSM&IDU | 324 | 30.82 | 100 | 12 |
| FSW | 1,978 | 4.08 | 81 | 20 |
| FSW&IDU | 62 | 61.72 | 38 | 12 |
| MCF | 61,596 | 1.06 | 653 | 8 |
| FPM | 454,074 | 0.45 | 2043 | 20 |

In order to provide a quantitative guide to controlling the epidemic we analyze the elements of the next-generation matrix (NGM) which give the case reproduction numbers (Diekmann et al., 2010) within and between groups. From the values of the coefficients in the model equations (Appendix), fitted to the time-series data (Figure 2), we obtain the elements of the NGM (Diekmann et al., 2010). The principal eigenvalue of the NGM is *R*_0_, the basic case reproduction number for the whole network; when *R*_0_ < 1, infection will be eliminated from the network (Diekmann *et al.*, 2010). Furthermore, on the approach to elimination, the smaller the value of *R*_0_, the smaller the number of people that will be infected. If a single infected case is introduced into one group then the elements of the NGM give the number of secondary cases that arise in each of the groups in the network and provide an elegantly simple method of investigating the impact of control measures, without resorting to specific numerical simulations and projections.

**Figure 2.**
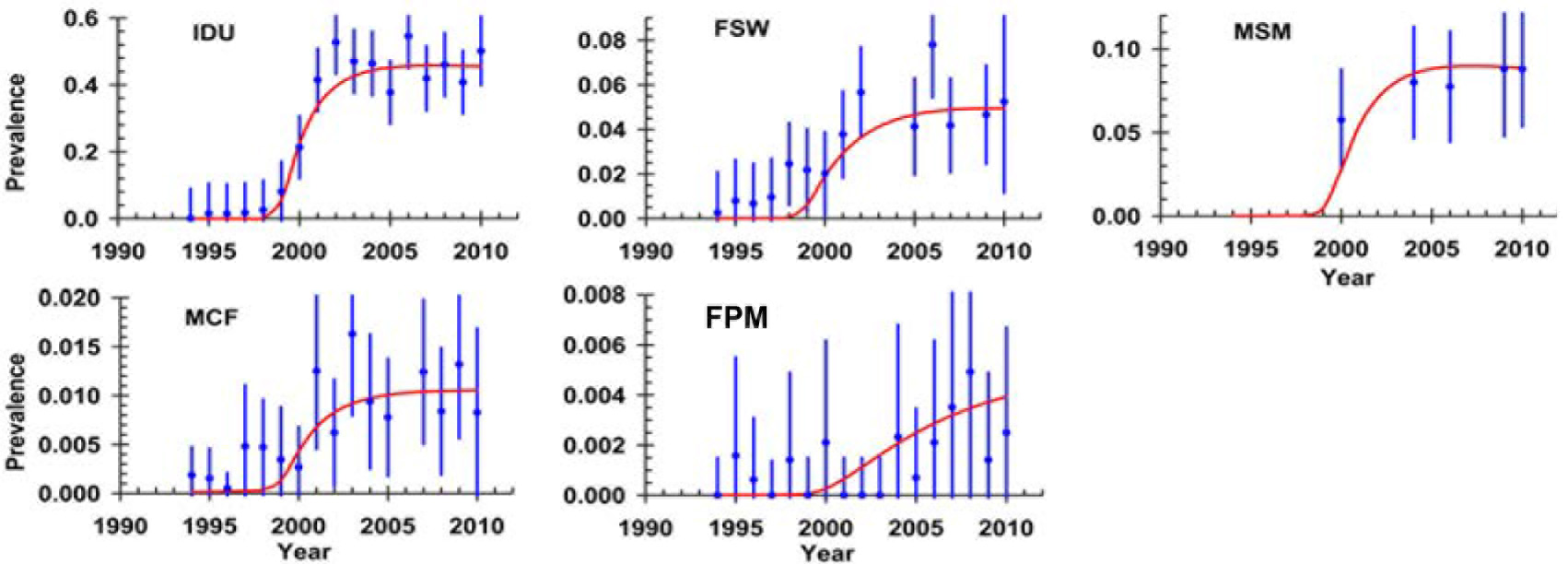
Trends in the prevalence of HIV over time for different risk groups in Can Tho province. IDU: Injection drug users; MSM: Men who have sex with men; FSW: Female sex workers; MCF: Male clients of FSWs; FPM: Female partners of MCF and other women at low risk.

## 3. Results

An earlier investigation of these data (Kato *et al.*, 2013) could not match the rapid rise in the prevalence of IDUs with the much slower rises in prevalence in other groups (Figure 2). To get a better fit to the data we assumed that infection was introduced initially among FSWs and then spread from them to IDUs. By setting the prevalence of HIV in the IDU group to zero in 1980, but allowing it to be non-zero in the other groups, we obtained the fit to the data shown in Table 1and Figure 2, which more accurately describes the spread of infection in all groups including IDUs.

Our first deduction from fitting the model to the time-series data is that the epidemic was probably introduced through female sex workers (FSW). From FSW it spread to injection drug users (IDU), who then became the key drivers of the epidemic. This conclusion is based on the observation that the model can accurately describe the epidemic in IDU only by assuming that HIV prevalence was zero in this group in 1980 and that infections in IDUs were introduced through the small group of FSWs who also inject drugs. The NGM gives *R*_0_ = 22 for the whole network, much larger than the value of *R*_0_ = 4.1 that would have been obtained by assuming random mixing among all the risk groups, assuming that they were all at equal risk, and fitting the model to time trends in the overall prevalence of HIV. The individual elements of the NGM give the values of *R*_0_ for transmission within and among population groups (Table 2). In Figure 3, values of *R*_0_ written in the circles apply within groups. Values of *R*_0_ written on the lines connecting groups give the number of secondary cases that arise from one primary case in the source group. Values of *R*_0_ written between the lines give the number of secondary cases arising in one primary group, via a linked secondary group.

**Table 2.** The next-generation matrix for the epidemic of HIV in Vietnam. For the whole system the eigenvalue of the dominant eigenvector gives *R*_0_ = 21.75. The last row gives the dominant eigenvector. The table gives the number of secondary cases in each group in a given row, as well as for all groups combined, for one primary case in each group in a given column in an otherwise fully susceptible population. Bullets mark cells that are identically zero. The elements of the matrix are calculated from the linearized equations, given in the Appendix, following Diekman et al. (2010).

|  | IDU | MSM | FSW | MCF | FPM | MSM<br>&IDU | FSW<br>&IDU |
| --- | --- | --- | --- | --- | --- | --- | --- |
| IDU | 19.27 | • | • | • | • | 19.27 | 19.27 |
| MSM | • | 4.09 | • | • | • | 4.09 | • |
| FSW | • | • | • | 0.058 | • | • | • |
| MCF | • | • | 77.27 | • | • | • | 77.27 |
| FPM | 0.007 | 0.001 | • | 0.010 | • | 0.000 | • |
| MSM&IDU | 2.001 | 1.18 | • | • | • | 3.329 | 2.00 |
| FSW&IDU | 0.469 | • | • | 0.003 | • | 0.469 | 0.93 |
| Total | 21.75 | 5.27 | 77.27 | 0.071 | • | 27.15 | 99.47 |
| Eigenvector | 0.989 | 0.025 | 0.000 | 0.088 | 0.000 | 0.111 | 0.025 |

If all of the connections between groups were broken, the IDU epidemic would still be self-sustaining in IDUs (*R*_0_ = 19). Similarly, the epidemic in FSW and MCF and the epidemic in MSM would each be self-sustaining but transmission in these groups is much easier to control because *R*_0_ is smaller (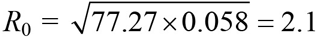 and 4.1, respectively). Because the IDU epidemic is linked to both MSM and FSW, control in the whole network will ultimately depend on control in IDUs.

Considering the links between pairs of groups (Figure 3), the most important are between IDU and MSM who are also IDU 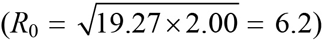 or FSW who are also IDU 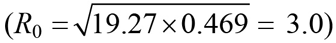. The loop connecting FSW and MCF is highly asymmetric: one case introduced in the FSW population will infect 77 MCF on average but each MCF infects only 0.058 FSW on average, over the life-time of an infected person. Thus, the number of secondary cases arising in FSW via MCF, over one complete cycle of transmission, is 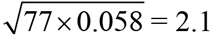 There are considerably fewer HIV-positive FSW than MCF (81 *versus* 653, Table 1) and each FSW has the potential to infect many more MCF over the ten years for which they will survive without treatment (77 *versus* 0.058, Table 2) so that, per person treated, interventions aimed at stopping transmission to and from FSW will be much more effective than interventions aimed at MCF.

**Figure 3.**
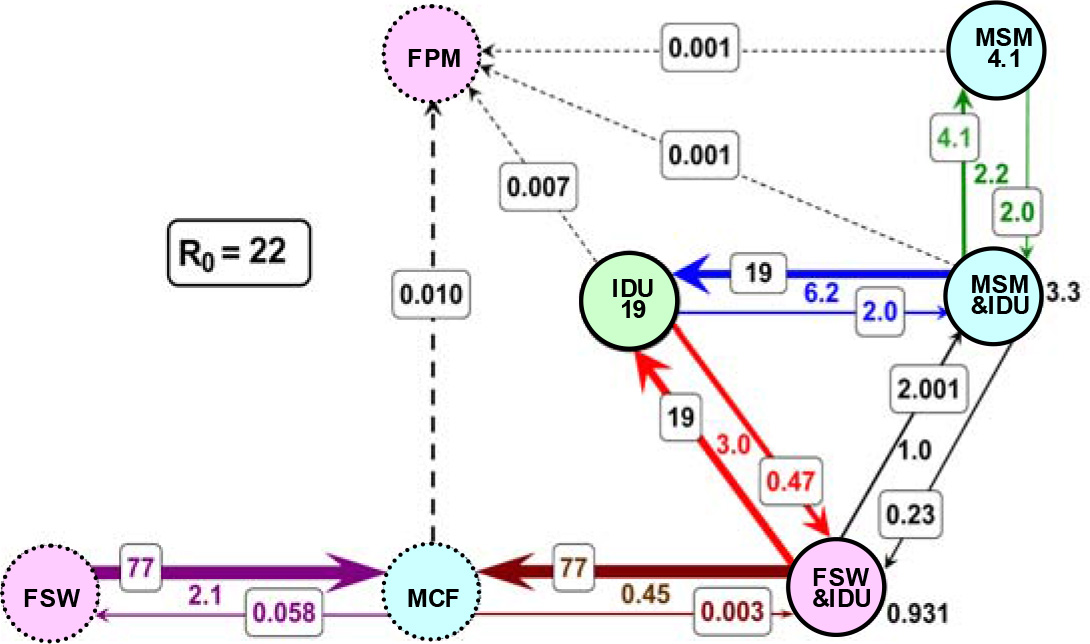
The network model with estimated transmission rates, which form the components of the next-generation matrix. Each number is the total number of secondary infections arising from one infected case in an otherwise susceptible population. Numbers in circles are for transmission within a population sub-group; numbers on lines are for transmission from one group to another; numbers between lines are the number of infections transmitted around a loop.

The epidemic of HIV in MSM (*R*_0_ = 2.2) should also be relatively easy to control. The prevention and treatment of infection in FPM is important in its own right and infected women should all have access to life-saving anti-retroviral therapy, but these women are assumed to be an epidemiological dead end, so control measures for these women will not affect transmission elsewhere in the network.

To choose the most effective control measures, the values of *R*_0_ need to be considered in relation to the efficacy of different interventions. To eliminate the epidemic within the IDU population requires *R*_0_ to be reduced by a factor of more than 19 (i.e. by more than 95%). Even with widespread use of antiretroviral therapy (ART)—for which the efficacy in preventing transmission from HIV-positive people to others has been estimated at 96% [12]— this is a challenge for ART when used as a single intervention and would demand very high levels of compliance and viral load suppression. However, combining ART with an effective needle and syringe exchange programme (a new needle and syringe carries zero risk of acquiring HIV) and opiate substitution therapy to reduce the use of injectable drugs, should be sufficient to achieve *R*_0_ < 1 in the IDU population (Montaner *et al.*, 2014). If, for example, half of the needle and syringe sharing involves clean needles and syringes then this would reduce *R*_0_ to about 10 and one would then only need a further 90% reduction through the use of ART to reduce *R*_0_ to less than 1; if clean needles and syringes were used in 95% of risky injection events then this alone would reduce *R*_0_ below 1 without the need for ART.

For each of the FSW and MSM populations it would be necessary to reduce transmission by a little more than 50% (more than 55% for *R*_0_ = 2.2 in MSM). For FSW, condom promotion would have a major impact and a ‘100% condom programme’ of the kind carried out in Thailand (Gouws *et al.*, 2006) should be sufficient to bring *R*_0_ below 1 for FSW and MCF, especially if supported by universal access to ART (MacArthur *et al.*, 2012). While consistent and correct condom use will completely stop the transmission of HIV, MSM may be reluctant to use condoms and condom promotion is generally found to be much less effective even under trial conditions (Williams, 2013). However, a programme of condom promotion combined with regular testing and universal access to ART should reduce *R*_0_ by more than the factor of 4.1 needed to control the epidemic among MSM.

Elimination of HIV from the whole network requires a combination of interventions against IDU, MSM, FSW and other groups. Further insights into the best combination of interventions that most effectively reduce *R*_0_ are provided by the NGM. Formally, elimination requires not only that *R*_0_ < 1 for each population group, but also for the network overall. Focusing on the key groups of IDU, MSM and FSW, let us assume that different numbers of each type can be removed from the pool of potentially infectious people. This could be achieved if people who were infected were immediately started on ART, if people used condoms in all sexual encounters or if IDUs were stopped injecting drugs through methadone maintenance programmes. We therefore consider the proportion of people of each type, involving different combinations of IDU, MSM and FSW, that are rendered non-infectious and removed from the model, and calculate the resulting value of *R*_0_ for the whole network. In what follows we use the word ‘treatment’ to indicate that people have been rendered non-infectious.

Figure 4A and 4B show, in principle, how to minimize *R*_0_ for the whole network by making the smallest number of people non-infectious. Notice that in each panel the lower contours of constant *R*_0_ are almost flat, so the best way to reduce *R*_0_ initially is by treating IDU alone (Figure 4A, vertical axis); there is little to be gained by treating members of other groups until a sufficiently large number of IDU have effectively been removed from the transmission network. The yellow dot in Figure 4A marks the position at which 2,400 IDU, but no MSM, are treated. Then, moving from the yellow dot, the best strategy is to treat both IDU and MSM until 2,700 IDU and 900 MSM are on treatment as indicated by the blue dot in Figure 4A. If one were only going to intervene with IDUs and MSM one would then continue along the line to the top right hand corner of Figure 4A when 3,100 IDUs and 1,150 MSM were on treatment. However, this would not eliminate transmission from the network as the epidemic in FSWs and MCFs is self-sustaining and the value of *R*_0_ for the whole network would be 2.2. The optimal strategy, after reaching the light blue point in Figure 4A or the corresponding light blue point in Figure 4B would be to increase the number of IDUs and MSM on treatment, keeping the proportion of each constant, but start treating FSWs following the curved line to the dark blue point in Figure 4B when *R*_0_ for the whole network would be reduced to 1. After that one would continue to the red dot in Figure 4B when all IDU, MSM and FSWs are rendered uninfectious and *R*_0_ for the whole network is reduced to 0.12.

The virtue of the NGM is that it gives an instant analytical guide to the question of where to focus interventions. To confirm the above results and also to explore the impact of different interventions through time demands a full dynamical simulation and this is illustrated in Figure 5.

**Figure 4.**
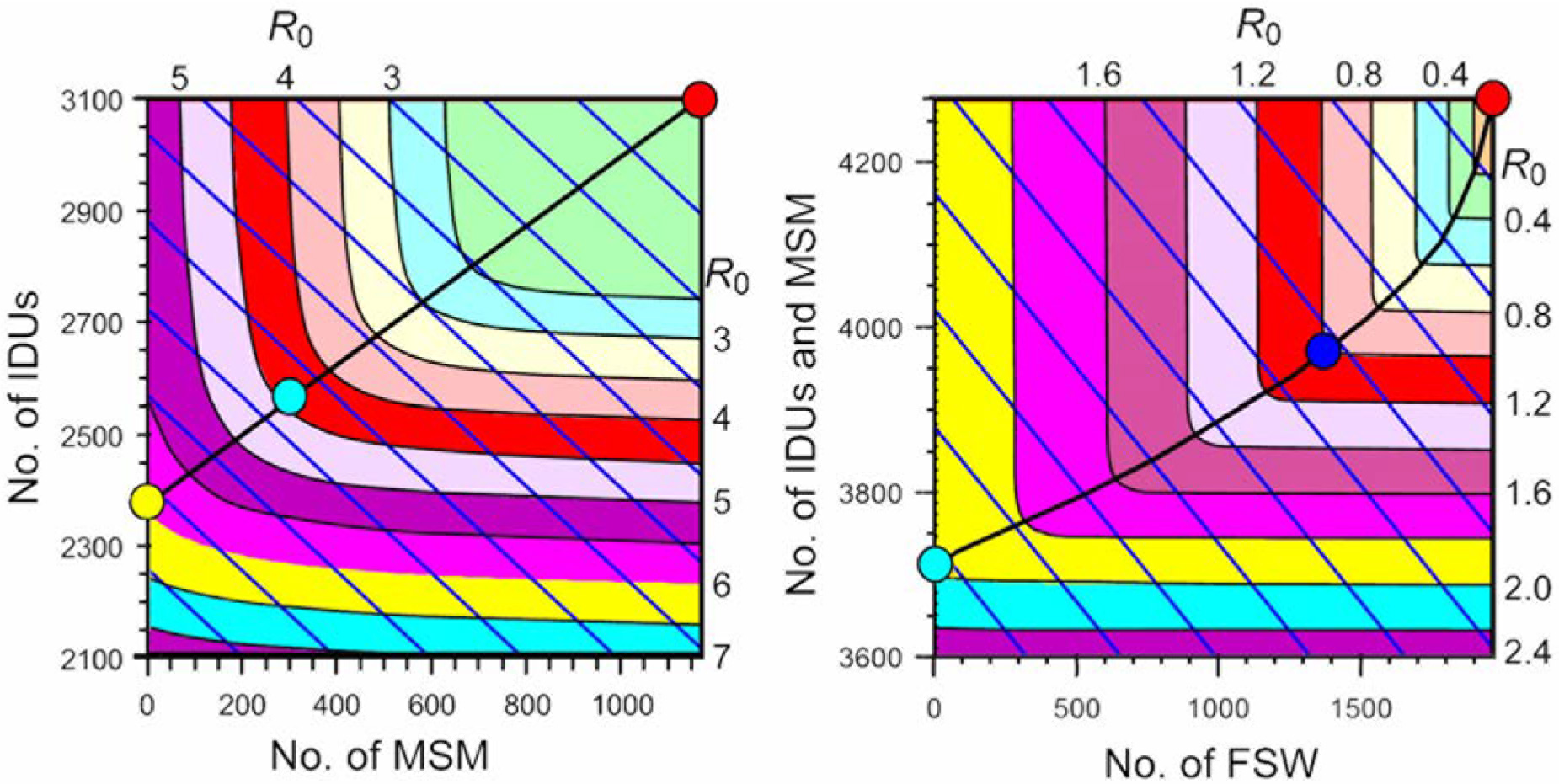
Surface plots of *R*_0_ as a function of the number of IDUs, MSM and FSWs who are rendered uninfectious either through treatment or prevention interventions. A: number of IDU plotted against the number of MSM who are rendered uninfectious. B: combined number of IDU and MSM plotted against the number of FSW who are rendered uninfectious. Shaded areas give contours of constant *R*_0_ for the values shown above and to the right of each plot. Diagonal lines in A indicate combinations of MSM and IDUs for fixed total numbers from 2,100 (bottom left) to 4,278 (top right); in B they indicate fixed total numbers of MSM and IDU (vertical axis) and FSW (horizontal axis) from 3,600 (bottom left) to 6,200 (top right). The lines running across each plot indicate the optimal combinations of IDUs, MSM and FSWs that minimize *R*_0_ for A: a fixed total number of IDUs and MSM and B: a fixed total number of IDUs, MSM and FSW.

Figure 5A shows the expected prevalence and incidence of HIV and AIDS-related mortality without any intervention in any group. This corresponds to the model fits given in Figure 2, projected forwards to 2050. Figure 5B shows what would happen in all five population groups if all IDU, but only IDU, were treated within one year of acquiring HIV infection so as to eliminate onward transmission. In Figure 5C to 5F the calculation is repeated for FSW, MCF, MSM and FPM.

Naturally, the treatment of people in any group reduces incidence, prevalence and mortality in that group. However, a comparison of Figure 5B with Figure 5C to 5F shows that only the treatment of IDU has a major impact on infections in all other population groups in the network. The secondary effect on MSM is most rapid, followed by FSW, MCF and FPM, as expected from the network structure shown in Figure 3. The treatment of FSW is also very beneficial for MCF (Figure 5C), but the reverse is not true (Figure 5D). Infection cannot be eliminated from the network by treating any one population group alone, though the treatment of IDU has the biggest overall impact. The benefits for other groups of treating MSM are small because MSM are weakly linked in the network (Figure 5E), except for the small proportion that also inject drugs (Figure 3). There are no benefits for other groups of treating FPM, because they are assumed not to transmit infection to anyone else (Figure 5F).

**Figure 5.**
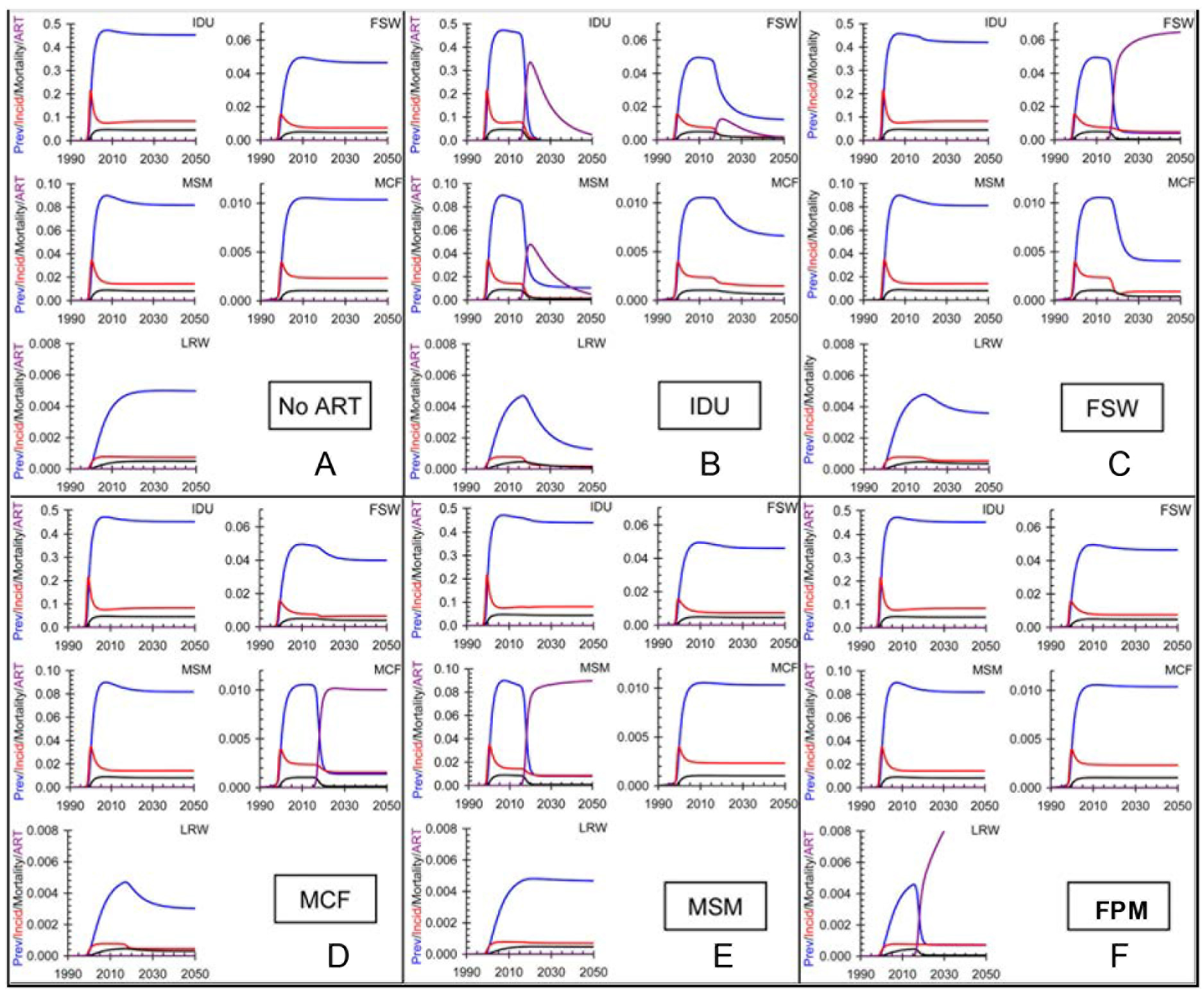
The boxes show the projected prevalence, incidence, mortality and ART coverage assuming that ART is provided to all those in the relevant population, as indicated in each box, with 50% coverage being reached in 2015 and full coverage in 2020 (see text for details). Blue lines: prevalence, red lines: annual incidence; black lines: annual mortality; purple lines: prevalence of people on ART.

## 4. Discussion

The better we understand the structure of transmission networks, the more effectively we can target efforts in infectious disease control. For concentrated epidemics of HIV/AIDS, the assumption of random mixing greatly underestimates the contribution of some population groups and greatly overestimates the contribution of others. This is illustrated here by the large difference in estimates of the basic case reproduction number of HIV estimated by assuming homogenous mixing (*R*_0_ = 4; data not shown) and derived from the structured social network (*R*_0_ = 22).

There is a cost to investigating the detailed structure of transmission networks, but the approach suggested here requires only data that are routinely collected during the spread of an epidemic, disaggregated for population groups that are likely to be exposed to infection at different rates, or transmit infection by different routes. As a further demonstration of heterogeneity, our reconstruction of the HIV/AIDS epidemic in Can Tho province shows that the infection was probably introduced first in FSW and MCF but then spread to IDU which became the main drivers of the epidemic. Ultimately the network generated most cases among the female partners of sex worker clients (Kato et al., 2013).

The structure and value of the elements in the next generation matrix give a guide to the key points at which control must be implemented and the degree of control that is needed to bring the epidemic to an end. Our analysis shows that IDU are the largest contributors (*R*_0_ = 19.3) to the overall case reproduction number (*R*_0_ = 22.0). The control of infection in IDU is the most effective way to reduce infections, not only in IDU, but across the whole network. To eliminate infection from the network altogether requires the reduction of *R*_0_ in IDU by a factor ≥ 19, but this must be achieved in combination with treatments for other population groups, especially FSW and MSM, so that *R*_0_ < 1 for every group separately and *R*_0_ < 1 for the network overall. Numerical simulations confirm these results, and show in detail how HIV incidence and prevalence and AIDS-related mortality can be expected to change through time. It should be noted that transmission from FSWs is mainly to MCFs who in turn infect their FPMs but these women do not infect others. Furthermore, if one treats IDUs this separates the MSM and FSW-MCF into two separate networks so that treating MSM has a greater impact on overall transmission than treating FSWs.

The optimal combination of prevention methods will depend on the group being targeted. For IDUs one would need a combination of opiate substitution therapy or methadone maintenance (MacArthur et al., 2012), access to clean needles and syringes (Aspinall et al., 2014), social support and ART (Cohen et al., 2011) as soon as people are found to be living with HIV. By combining these interventions one would get significant synergies. If, as is the case in Vietnam, methadone maintenance programmes require daily attendance by patients with a medical doctor present at all times, this would provide an ideal setting for the provision of anti retroviral drugs combined with testing viral loads to ensure compliance. For FSWs in brothels a condom programme of the kind rolled out in Thailand should have a significant effect on transmission (Park et al., 2010) and this could be combined with routine testing of FSWs for HIV, if they are previously HIV-negative, and viral loads, if they are already infected with HIV.

A full uncertainty analysis could be done using a Bayesian approach to constrain the parameters where the data are less certain. This analysis suggests that to reduce *R*_0_ most efficiently one should focus first on IDUs, then on MSM and finally on FSWs. Because the data on MSM are very sparse it is possible that the initial rate of increase, and hence the value of *R*_0_ for MSM is greater than estimated here. One might wish to explore further the effect of uncertainty in the trend data for MSM on the overall conclusions.

Although the NGM constructed from routine data gives insights into HIV epidemiology and control quickly and relatively easily, it is not the last word in analysis. For instance, genotyping studies could help to confirm or refute our deduction that the epidemic in IDU was first introduced by FSW. We have also assumed that the population of Can Tho province is affected by a single strain of HIV even though, in other settings, MSM may be infected by different strains of HIV from FSW and MCF giving rise to separate epidemics (Dennis et al., 2014). Because our method of analysing epidemic spread and control requires only routine data, it could potentially be applied to HIV infection and other communicable diseases in different populations. However, it would be instructive and prudent to carry out an analysis of HIV strain variation in HIV infections in any other population to which this method of network analysis might be applied.

Here we are concerned to demonstrate the information that can be gained from a detailed analysis of the structure of the NGM as a guide to the choice of interventions and to facilitate a complete analysis of future projections, estimates of impact, and costs and cost effectiveness of different interventions. In this, as in all public health data, there is uncertainty in the data, and hence in the fitted curves and corresponding parameter estimates and these should be used to add uncertainty estimates to the various fitted parameters, estimates of *R*_0_ and future projections.

It is, of course, important to bear in mind that many different objective functions can be chosen; here we have chosen to minimize *R*_0_, others might wish to minimize the total cost of the intervention, the cost per infection averted or per life saved, for example, and each of these would lead to a different optimal strategy. But all these options could be explored with the network analysis we have described here.

In each particular setting, it will be necessary to identify the relevant risk groups, decide on the optimal combination of interventions for each group taking into account both the efficacy and the cost of each component intervention, and then plan the roll-out of the control programme accordingly. Our intention here is not to provide a definitive answer to the best way to manage HIV in Can Tho and we have only considered what would happen if one were to effectively reduce the number of people of each type that could potentially contribute to HIV transmission. In a more detailed analysis, managers of control programmes might, for example, note that male circumcision provides a 60% reduction in transmission from women to men but none from men to women. They could then calculate the reduction in transmission from men to women. What we are suggesting is that, having fitted a model to a complex epidemic involving disparate population types, as is the case in Can Tho, one can use the ideas behind the Type Reproduction number and the Target Reproduction Number to explore the impact of different intervention strategies, directly from the next generation matrix, based on routinely collected data, and without having to run and examine the full dynamical model over time.

Eventually, a full dynamical model will have to be used to evaluate the long term impact, on the HIV prevalence, incidence and mortality as well as the cost and cost-effectiveness of alternative interventions An analysis of the kind presented in this paper should, however, provide a useful and informative starting point for thinking about the best way to control HIV.

## Appendix: Model equations

The model used in this analysis is illustrated schematically in Figure 1. The structure, and in particular, the overlapping groups and the links between pairs of groups, was arrived at after extensive discussions with field workers supporting each of the risk groups in Vietnam (Kato et al., 2013). A critical risk group consists of female sex workers who also inject drugs and form a bridging population between those who are at risk through heterosexual transmission and those who are at risk through the sharing of contaminated needles and syringes. Most of these women are primarily injecting drug users who do sex work to raise money to buy drugs.

The number of people in various classes is 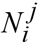 where *j* defines a subset of people in class *i*. If there is no superscript the number refers to the whole class defined by the subscript. For example, *N*_*m*_ is the total number of MSM; 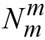 is the number of MSM who are only MSM; 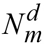 is the number of MSM who are also IDUs etc. Index *s* refers to FSW, *c* to the male clients of FSWs, *w* to the female partners of male clients of FSWs. For each class 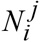 the number of infected people is 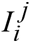 and the number of susceptible people is 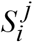. We let

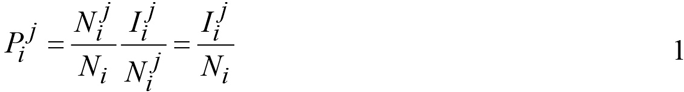

so that 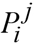 gives the proportion of those that are in class *i* that are also in class *j* times the prevalence in those that are in class *i* and *j*. For example,

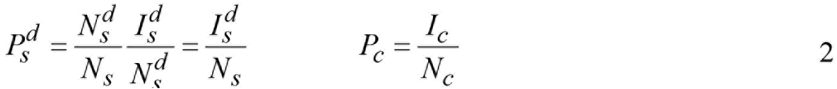

The lower subscript determines the group, which may be *d*, *m*, *s*, *c*, *w* and the upper subscript the route of transmission for that group, which may be *d*, *m* or *s*. The equations for the model are then

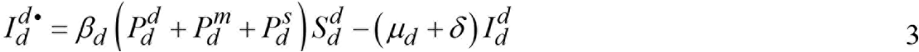

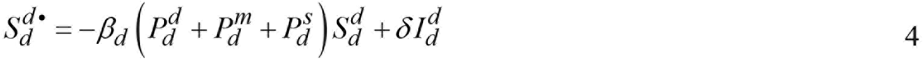

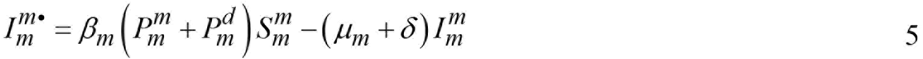

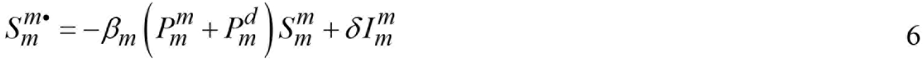

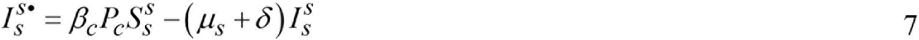

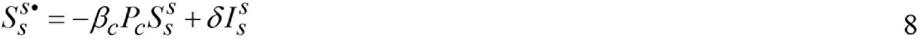

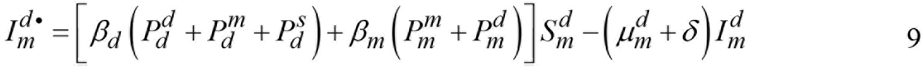

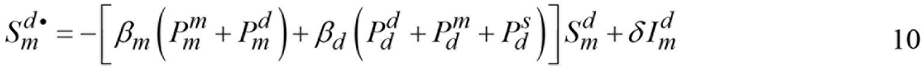

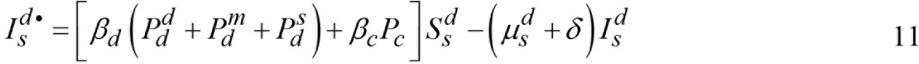

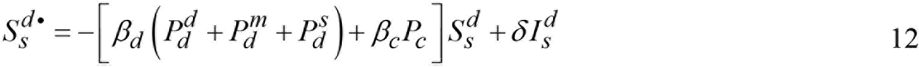

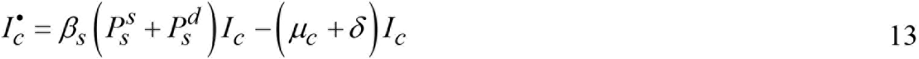

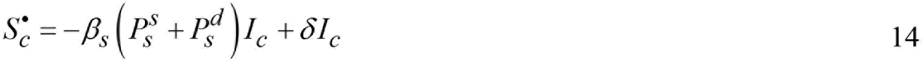

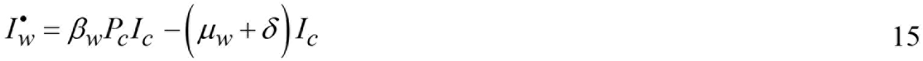

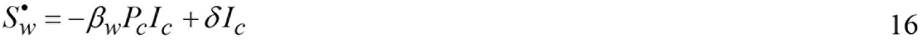

The next generation matrix is given in Table A1.

**Table A1.** The next generation matrix corresponding to the model specified in Equations 1 to 16. *δ* without a subscript refers to the background mortality which we take to be the same for all groups and we set this to 0.02/year corresponding to a life expectancy of 50 years. The chance of being infected through drug use is independent of whether or not that person is also MSM or FSW. In practice MSM who also use drugs may be more likely to be infected by other MSM rather than FSW who also use drugs

| | $d$ | $m$ | $s$ | $dm$ | $ds$ | $c$ | $w$ |
| --- | --- | --- | --- | --- | --- | --- | --- |
| $d$ | $\frac{\beta_d P_d^d}{\mu_d + \delta}$ | | | $\frac{\beta_d P_d^m}{\mu_d + \delta}$ | $\frac{\beta_d P_d^s}{\mu_d + \delta}$ | | |
| $m$ | | $\frac{\beta_m P_m^m}{\mu_m + \delta}$ | | $\frac{\beta_m P_m^d}{\mu_m + \delta}$ | | | |
| $s$ | | | | | | $\frac{\beta_c P_c}{\mu_s + \delta}$ | |
| $dm$ | $\frac{\beta_d P_d^d}{\mu_d^m + \delta}$ | $\frac{\beta_m P_m^m}{\mu_d^m + \delta}$ | | $\frac{(\beta_d + \beta_m) P_d^m}{\mu_d^m + \delta}$ | $\frac{\beta_d P_d^s}{\mu_d^m + \delta}$ | | |
| $ds$ | $\frac{\beta_d P_d^d}{\mu_d^s + \delta}$ | | | $\frac{\beta_d P_d^m}{\mu_d^s + \delta}$ | $\frac{\beta_d P_d^s}{\mu_d^s + \delta}$ | $\frac{\beta_c P_c}{\mu_d^s + \delta}$ | |
| $c$ | | | $\frac{\beta_s P_s^s}{\mu_c + \delta}$ | | $\frac{\beta_s P_s^d}{\mu_c + \delta}$ | | |
| $w$ | | | | | | | $\frac{\beta_w}{\mu_w + \delta}$ |

In order to allow for heterogeneity in risk, which determines the steady state prevalence of infection, we assume that the rate of transmission, *β* in these equations are multiplied by a corresponding Gaussian term so that

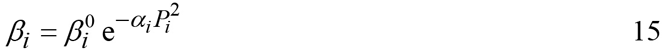

so that 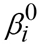 is the rate of transmission in group *i* at the start of the epidemic when the prevalence is close to zero and the rate of transmission fall as the prevalence *P*_*i*_ = *I*_*i*_/*N*_*i*_.rises. At the start of the epidemic the prevalence is low but those at highest risk will be infected first.

As prevalence rises, those that are not yet infected will be at lower risk and the average value of the transmission parameter will decrease as the prevalence of infection increases. Previous studies have assumed an exponential relationship or a step-function. In the former case the solutions tend to be unstable as the risk of infection drops rapidly and prevalence increases rapidly as prevalence declines. In the latter case one is dividing the population into those at a certain fixed risk and those at no risk. The prevalence data can be fitted equally well under either of these two extreme assumptions and there is, unfortunately, no direct evidence to determine the rate at which transmission falls as prevalence rises. This is an area that warrants further investigation (Williams et al., 2015).

The model fits are given in Figure 2 and the parameter values for the fits are given in Table A2. The time for which people remain in a risk group and the size of each risk group were obtained from field workers supporting each of the risk groups in Vietnam (Kato et al., 2013).

**Table A2.** Best fit values of the transmission parameters, estimated durations within each risk group and estimated size of each risk group. The loss rate is the rate at which people leave each group. The AIDS related mortality is 0.1/year.

| Transmission/yr |  | Loss rate/yr |  | Group size |  |
| --- | --- | --- | --- | --- | --- |
| $\beta_d$ | 2.51 | $\mu_d$ | 0.084 | $N_d$ | 3,698 |
| $\beta_m$ | 0.08 | $\mu_m$ | 0.050 | $N_m$ | 1,183 |
| $\beta_c$ | 0.34 | $\mu_c$ | 0.100 | $N_s$ | 2,011 |
| $\beta_s$ | 0.24 | $\mu_s$ | 0.125 | $N_{md}$ | 384 |
| $\beta_w$ | 0.00 | $\mu_w$ | 0.050 | $N_{sd}$ | 90 |
| | | | | $N_c$ | 62,000 |
| | | | | $N_w$ | 455,141 |

## References

Aspinall, E.J., Nambiar, D., Goldberg, D.J., Hickman, M., Weir, A., Van Velzen, E., Palmateer, N., Doyle, J.S., Hellard, M.E., Hutchinson, S.J., 2014. Are needle and syringe programmes associated with a reduction in HIV transmission among people who inject drugs: a systematic review and meta-analysis. International journal of epidemiology 43, 235–248.

Brenner, B.G., Wainberg, M.A., 2013. Future of Phylogeny in HIV Prevention. Journal of Acquired Immune Deficiency Sydnromes 63, S248–254.

Cohen, M.S., Chen, Y.Q., McCauley, M., Gamble, T., Hosseinipour, M.C., Kumarasamy, N., Hakim, J.G., Kumwenda, J., Grinsztejn, B., Pilotto, J.H., Godbole, S.V., Mehendale, S., Chariyalertsak, S., Santos, B.R., Mayer, K.H., Hoffman, I.F., Eshleman, S.H., Piwowar-Manning, E., Wang, L., Makhema, J., Mills, L.A., de Bruyn, G., Sanne, I., Eron, J., Gallant, J., Havlir, D., Swindells, S., Ribaudo, H., Elharrar, V., Burns, D., Taha, T.E., Nielsen-Saines, K., Celentano, D., Essex, M., Fleming, T.R., 2011. Prevention of HIV-1 infection with early antiretroviral therapy. New England Journal of Medicine 365, 493–505.

Cohen, M.S., McCauley, M., Gamble, T.R., 2012. HIV treatment as prevention and HPTN 052. Current opinion in HIV and AIDS 7, 99–105.

Dennis, A.M., Herbeck, J.T., Brown, A.L., Kellam, P., de Oliveira, T., Pillay, D., Fraser, C., Cohen, M.S., 2014. Phylogenetic Studies of Transmission Dynamics in Generalized HIV Epidemics: An Essential Tool Where the Burden is Greatest? Journal of Acquired Immune Deficiency Syndromes 67, 181–195.

Diekmann, O., Dietz, K., Heesterbeek, J.A., 1991. The basic reproduction ratio for sexually transmitted diseases: I. Theoretical considerations. Math Biosci 107, 325–339.

Diekmann, O., Heesterbeek, J.A., Metz, J.A., 1990. On the definition and the computation of the basic reproduction ratio R0 in models for infectious diseases in heterogeneous populations. Journal of mathematical biology 28, 365–382.

Diekmann, O., Heesterbeek, J.A.P., Roberts, M.G., 2010. The construction of next-generation matrices for compartmental epidemic models. Journal of the Royal Society Interface 7, 873–885.

Gouws, E., Cuchi, P., 2012. Focusing the HIV response through estimating the major modes of HIV transmission: a multi-country analysis. Sexually Transmitted Infections 88 Suppl 2, i76–85.

Gouws, E., White, P.J., Stover, J., Brown, T., 2006. Short term estimates of adult HIV incidence by mode of transmission: Kenya and Thailand as examples. Sexually Transmitted Infections 82, S51–55.

Grabowski, M.K., Lessler, J., Redd, A.D., Kagaayi, J., Laeyendecker, O., Ndyanabo, A., Nelson, M.I., Cummings, D.A.T., Bwanika, J.B., Mueller, A.C., Reynolds, S.J., Munshaw, S., Ray, S.C., Lutalo, T., Manucci, J., Tobian, A.A.R., Chang, L.W., Beyrer, C., Jennings, J.M., Nalugoda, F., Serwadda, D., Wawer, M.J., Quinn, T.C., Gray, R.H., the Rakai Health Sciences, P., 2014. The Role of Viral Introductions in Sustaining Community-Based HIV Epidemics in Rural Uganda: Evidence from Spatial Clustering, Phylogenetics, and Egocentric Transmission Models. PLOS Medicine 11, e1001610.

Heesterbeek, H., Anderson, R., Andreasen, V., Bansal, S., De Angelis, D., Dye, C., Eames, K., Edmunds, J., Frost, S., Funk, S., Hollingsworth, D., House, T., Isham, V., Klepac, P., Lessler, J., Lloyd-Smith, J., Metcalf, J., Mollison, D., Pellis, L., Pulliam, J., Roberts, M., Viboud, C., 2015. Modeling infectious disease dynamics in the complex landscape of global health. Science 347, aaa4339–aaa4339.

Heesterbeek, J.A., 2002. A brief history of R0 and a recipe for its calculation. Acta biotheoretica 50, 189–204.

Heesterbeek, J.A., Roberts, M.G., 2007. The type-reproduction number T in models for infectious disease control. Math Biosci 206, 3–10.

Helleringer, S., Kohler, H.P., Chimbiri, A., Chatonda, P., Mkandawire, J., 2009. The Likoma Network Study: Context, data collection, and initial results. Demographic research 21, 427–468.

Kato, M., Granich, R., Bui, D.D., Tran, H.V., Nadol, P., Jacka, D., Sabin, K., Suthar, A.B., Mesquita, F., Lo, Y.R., Williams, B., 2013. The potential impact of expanding antiretroviral therapy and combination prevention in Vietnam: towards elimination of HIV transmission. Journal of Acquired Immune Deficiency Syndromes 63, e142–149.

Leventhal, G.E., Kouyos, R., Stadler, T., Wyl, V., Yerly, S., Boni, J., Cellerai, C., Klimkait, T., Gunthard, H.F., Bonhoeffer, S., 2012. Inferring epidemic contact structure from phylogenetic trees. PLoS Computational Biology 8, e1002413.

Lurie, M., Williams, B.G., Zuma, K., Mkaya-Mwamburi, D., Garnett, G.P., Sweat, M.D., Gittelsohn, J., Abdool Karim, S., 2003. Who infects whom? HIV-1 concordance and discordance among migrant and non-migrant couples in South Africa. Aids 17, 2245–2252.

MacArthur, G.J., Minozzi, S., Martin, N., Vickerman, P., Deren, S., Bruneau, J., Degenhardt, L., Hickman, M., 2012. Opiate substitution treatment and HIV transmission in people who inject drugs: systematic review and meta-analysis. British medical journal 345, e5945.

Montaner, J.S.G., Lima, V.D., Richard Harrigan, P.R., Lourenc, L., Yio, B., Nosyk, B., Wood, E., Kerr, T., Shannon, K., Moore, D., Hogg, R.S., Barrios, R., Gilbert, M., Krajden, M., Gustafson, R., Daly, P., Kendall, P., 2014. Expansion of HAART coverage Is associated with sustained decreases in HIV/AIDS morbidity, mortality and HIV transmission: The ‘HIV Treatment as Prevention’ experience in a Canadian setting. PLOS One 9(2): e87872.

Park, L.S., Siraprapasiri, T., Peerapatanapokin, W., Manne, J., Niccolai, L., Kunanusont, C., 2010. HIV transmission rates in Thailand: evidence of HIV prevention and transmission decline. Journal of Acquired Immune Deficiency Syndromes and Human Retrovirology 54, 430–436.

Roberts, M.G., Heesterbeek, J.A., 2003. A new method for estimating the effort required to control an infectious disease. Proceedings. Biological sciences / The Royal Society 270, 1359–1364.

Roberts, M.G., Heesterbeek, J.A., 2007. Model-consistent estimation of the basic reproduction number from the incidence of an emerging infection. Journal of mathematical biology 55, 803–816.

Roberts, M.G., Heesterbeek, J.A., 2012. Characterizing the next-generation matrix and basic reproduction number in ecological epidemiology. Journal of mathematical biology.

Sattenspiel, L., Simon, C.P., 1988. The spread and persistence of infectious diseases in structured populations. Math Biosci 90, 341–366.

Shuai, Z., Heesterbeek, J.A.P., van den Driessche, P., 2013. Extending the type reproduction number to infectious disease control targeting contacts between types. Journal of mathematical biology 67, 1067–1082.

Stadler, T., Kouyos, R., von Wyl, V., Yerly, S., Boni, J., Burgisser, P., Klimkait, T., Joos, B., Rieder, P., Xie, D., Gunthard, H.F., Drummond, A.J., Bonhoeffer, S., 2012. Estimating the basic reproductive number from viral sequence data. Molecular biology and evolution 29, 347–357.

Williams, B.G., 2013. Combination prevention for the elimination of HIV, arXiv.

Williams, B.G., Gouws, E., Somse, P., Mmelesi, M., Lwamba, C., Chikoko, T., Fazito, E., Turay, M., Kiwango, E., Chikukwa, P., Damisoni, H., Gboun, M., 2015. Epidemiological Trends for HIV in Southern Africa: Implications for Reaching the Elimination Targets. Current HIV/AIDS reports 12, 1–11.

